# Reconstructing genomes of carbon monoxide oxidisers in volcanic deposits including members of the class Ktedonobacteria

**DOI:** 10.1101/2020.10.29.361295

**Authors:** Marcela Hernández, Blanca Vera-Gargallo, Marcela Calabi-Floody, Gary M King, Ralf Conrad, Christoph C. Tebbe

## Abstract

Microorganisms can potentially colonize volcanic rocks using the chemical energy in reduced gases such as methane, hydrogen (H_2_) and carbon monoxide (CO). In this study, we analysed soil metagenomes from Chilean volcanic soils, representing three different successional stages with ages of 380, 269 and 63 years, respectively. A total of 19 metagenome-assembled genomes (MAGs) were retrieved from all stages with a higher number observed in the youngest soil (1640: 2 MAGs, 1751: 1 MAG, 1957: 16 MAGs). Genomic similarity indices showed that several MAGs had amino-acid identity (AAI) values >50% to the phyla Actinobacteria, Acidobacteria, Gemmatimonadetes, Proteobacteria and Chloroflexi. Three MAGs from the youngest site (1957) belonged to the class Ktedonobacteria (Chloroflexi). Complete cellular functions of all the MAGs were characterised, including carbon fixation, terpenoid backbone biosynthesis, formate oxidation and CO oxidation. All 19 environmental genomes contained at least one gene encoding a putative carbon monoxide dehydrogenase (CODH). Three MAGs had form I *coxL* operon (encoding the large subunit CO-dehydrogenase). One of these MAGs (MAG-1957-2.1, Ktedonobacterales) was highly abundant in the youngest soil. MAG-1957-2.1 also contained genes encoding a [NiFe]-hydrogenase and *hyp* genes encoding accessory enzymes and proteins. Little is known about the Ktedonobacterales through cultivated isolates, but some species can utilize H_2_ and CO for growth. Our results strongly suggest that the remote volcanic sites in Chile represent a natural habitat for Ktedonobacteria and they may use reduced gases for growth.

## 1. Introduction

Volcanic eruptions provide a model for understanding soil-forming processes and the roles of pioneer bacteria during early biotic colonization. Recently, it has been demonstrated that the structure of microbial communities can play a key role in the direction of plant community succession pathways [1]. This is due in part to bacterial contributions to weathering of volcanic rocks, which releases nutrients resulting in some of the most fertile soils in the world.

After lava and other volcanic deposits (i.e. ash and tephra) cool sufficiently, mineral surface areas become accessible for microbial colonization [2–5]. In fact, microbes, and especially bacteria, are among the first colonizers of volcanic deposits and thereby initiate soil formation during the early stages of terrestrial ecosystem development [6–12]. While methane (CH_4_), hydrogen sulfide (H_2_S), hydrogen (H_2_), and carbon monoxide (CO) have been proposed to promote bacterial colonization and support microbial life in these organic-carbon deficient environments [13,14], the actual carbon and energy sources of the first colonizers remain elusive, but likely include a range of endogenous and exogenous sources, including reduced minerals and gases. The microbial ability to utilize these substrates for growth, and thereby initiate the formation of soil organic material, depends on specialized enzymes which may not be prevalent in many different microbial groups, thus representing a limited phylogenetic distribution. As a consequence, this could constrain the composition of pioneering microbial communities.

CO is a potential source of carbon and energy for microbes pioneering the colonization of volcanic substrates. CO utilization under oxic conditions requires a molybdenum-dependent carbon monoxide dehydrogenase (Mo-CODH), which catalyses the oxidation of CO to CO_2_ [15]. Surveys of genome databases (e.g., Integrated Microbial Genomes) reveal that Mo-CODHs occur in Proteobacteria, Actinobacteria, Firmicutes, Chloroflexi, Bacteroidetes, Deinococcus-Thermus, Halobacteria, and Sulfolobales among others. They were originally described as inducible enzymes and have subsequently been shown to be up-regulated by carbon limitation during growth of Actinobacteria (e.g., *Rhodococcus* and *Mycobacterium smegmatis* [15–17], and Chloroflexi (e.g., *Thermogemmatispora* and *Thermomicrobium* [18]). Mo-CODH has been previously targeted in molecular ecological studies of CO oxidizers in volcanic systems and other extreme environments [15]. These studies have revealed changes in community composition with the age and developmental status of individual sites [7,15]. The results suggest that CO-oxidizing communities are not static, but that they change in response to changing environmental conditions, and possibly affect the direction of changes.

In a preceding study, we identified the microbial communities involved in volcanic soil formation in different sites on Llaima Volcano (Chile). The bacterial communities of soils from 3 sites affected by lava deposition in 1640, 1751 and 1957 were analysed using 16S rRNA gene amplicon sequencing [19] and it was demonstrated that microbial diversity increased with the age of the soil deposits. Interestingly, bacterial phylotypes of the poorly studied Ktedonobacterales were among the predominant community members in the 1957 soil, representing 37% of all OTUs, as compared with 18% in the 1751 and 7% in the 1640 soils. Thus, we suspected that bacteria of this order could be instrumental for the initiation of soil formation, paving the way for soil organic carbon formation and preparing a substrate for microbial colonisation and plant growth. Some already cultivated Ktedonobacterales were found to be carboxydotrophs and hydrogenotrophs (*i.e.* carbon monoxide (CO) and hydrogen (H_2_) oxidisers/consumers) [14]. Thus, this leads to the hypothesis that CO and H_2_ are important carbon and energy sources for early stages of microbes colonizing the Llaima Volcano soil. However, the ecophysiology of the few bacterial isolates assigned to Ktedonobacterales limits predictions about metabolic functions based on 16S rRNA gene sequences alone.

Therefore, in this study a metagenomic approach was chosen to identify microbial traits associated with early stages of colonization and soil formation in a volcanic ecosystem. In particular, the presence of functional genes implicated in CO-oxidation (*coxL* genes) and H_2_-oxidation (*hyd* and *hyp* genes) was assessed from metagenome-assembled genomes (MAGs). These MAGs were retrieved from volcanic soils of different ages, representing sites 1640, 1751 and 1957. During this period, the soil formation evolved as indicated by their different levels of soil organic matter ranging from 65.33 ± 2.31 in the most recent soil (1957) to 9.33 % in both medium (1751) and oldest soils (1640) [20].

Especially the most recent soil (youngest soil) was suspected to reveal microbial adaptations to the challenging environmental conditions and thus to unveil the metabolic processes which initiate microbial colonisations. Therefore, functional metabolic modules annotated in the environmental genomes were analysed, with a main focus on the poorly characterised class of the Ktedonobacteria (Chloroflexi). Three Ktedonobacteria MAGs were obtained and all contained genes encoding CO and H_2_ oxidation. Additional MAGs from other phyla were also found to contain these genes. Our study advances the understanding of the ecology of Ktedonobacteria and their potential to act as early colonizers in volcanic soils.

## 2. Materials and Methods

### 2.1 Sequencing

The DNA from the volcanic soils used in our study had been previously extracted [20]. The soil physico-chemical characteristics have been published [20] showing a pH of 5.6 in both medium and oldest soil and 4.7 in the youngest soil, and nitrogen (mg/kg) of 25 (1640), 26 (1751) and 36.33 (1957). Briefly, the soil samples originating from three different sites of different ages according to the latest lava eruption (1640, 1751, 1957, map in [20]). A total of nine samples (triplicate per site) were sequenced on an Illumina MiSeq at the Max-Planck-Genome Centre, Köln, Germany. The metagenome was analysed on a high-performance computer using 650 GB RAM and 64 cores at the Thünen Institute of Biodiversity, Braunschweig, Germany.

### 2.2. Quality control

The sequence reads were checked using FastQC version 0.11.8 [21]. Low quality reads were discarded using BBDuk version 38.68, quality-trimming to Q15 using the Phred algorithm [22]. A schematic overview of the steps and programs used are shown in Figure 1.

**Figure 1.**
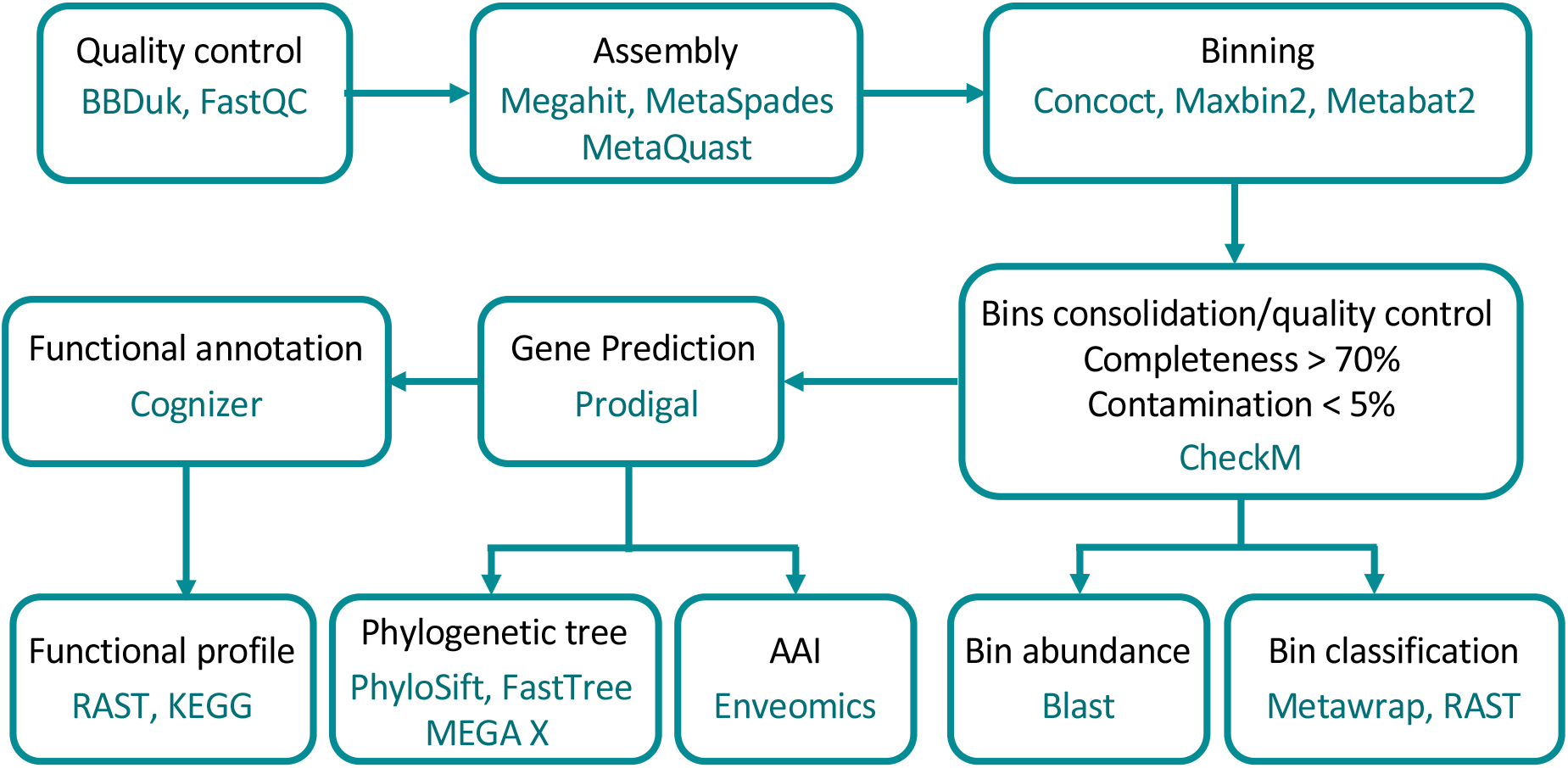
Workflow of the metagenome analysis and programs used in the present study. Binning was run by using MetaWrap package. AAI: Amino Acid Identity.

### 2.3. Metagenome assembly and binning

All trimmed Illumina reads were merged into longer contiguous sequences (scaffolds) using *de novo* assemblers Megahit version 1.2.8 [23] with k-mers 21,29,39,59,79,99,119,141 and MetaSPAdes (SPAdes for co-assembly) version 3.13.1 [24,25] with k-mers 21,31,41,51,61,71,81. Triplicate samples were co-assembled in order to improve the assembly of low-abundance organisms. Assembly quality was checked with MetaQuast version 5.0.2 [26] showing that the best quality was obtained with MetaSPAdes for our samples (data not shown). Downstream analysis was carried out using the scaffolds retrieved from MetaSPAdes. Krona charts [27] were recovered from MetaQuast runs to identify taxonomic profiles. Downstream binning analysis was performed with two sets of scaffolds: full size scaffolds and scaffolds larger than 1000 bp.

Metagenomic binning of the assembled scaffolds was carried out with the metaWRAP version 1.2.1 pipeline [28], which binning module employs three binning software programs: MaxBin2 [29], metaBAT2 [30], and CONCOCT [31]. Completion and contamination metrics of the extracted bins were estimated using CheckM [32]. The resulting bins were collectively processed to produce consolidated metagenome-assembled genomes (MAGs) using the bin_refinement module (criterion: completeness > 70%; contamination < 5%). Both sets of MAGs (18 from scaffolds larger than 1000 bp and 17 from full size scaffolds) were aggregated, visualized with VizBin [33] and then dereplicated using dRep [34]. Only the highest scoring MAG from each secondary cluster was retained in the dereplicated set. The abundance of each MAG in the different sites was calculated using BLASTN version 2.5.0+ [35] keeping only hits with >95% identity and e-value 1e-5 for the analysis [36]. A final heatmap was constructed using the function heatmap.2 from the gplots package version 3.0.4 [37] in R version 4.0.2 (https://www.r-project.org).

### 2.4. Functional annotation

The open reading frames (ORFs) in all scaffolds of each MAG were predicted using Prodigal (v2.6.3) [38]. Functions were annotated using Cognizer [39] and KEGG annotation framework [40]. The annotations of the predicted proteins from the Kyoto Encyclopedia of Genes and Genomes (KEGG) were used to confirm protein functional assignment and identify pathways. Complete pathways were identified using KEGG BRITE pathway mapping [40]. Aerobic carbon-monoxide dehydrogenases and hydrogen dehydrogenase were also identified using KEGG ortholog annotations; CODH was further distinguished as form I and form II (putative CODH) based on active site motifs present in *coxL* genes (e.g., [41]).

### 2.5. Phylogenomic analysis

Taxonomic classification of MAGs was performed using the classify_bins module from metaWRAP which relies on NCBI_nt database. MAGs were also screened using the RAST Server (Rapid Annotations using Subsystems Technology; [42,43]), which also allowed to retrieve information regarding close relative genomes in order to construct the phylogenetic tree.

To estimate intergenomic similarity, amino-acid comparisons between MAGs and their closest relative genomes present in the databases were calculated based on reciprocal best hits (two-way AAI) using the enveomics collection (http://enve-omics.gatech.edu/ [44].

The phylogenetic affiliation of MAGs was determined by constructing a genomic tree using FastTree version 2.1.11 [45]. Reference genomes were manually downloaded from the National Center for Biotechnology Information (NCBI) Refseq database (Table S1). Conserved genes from the extracted bins and the reference genomes were concatenated using Phylosift version 1.0.1 [46].

Phylogenetic analysis of the large sub-unit CO dehydrogenase gene (*coxL*) using the Maximum Likelihood method with a JTT matrix-based model [47] was performed. Bootstrap values (100 replicates) are shown where support ≥ 70 percent. The scale bar indicates substitutions per site. All gapped positions were deleted resulting in 420 positions in the final dataset. Evolutionary analyses were conducted in MEGA X [48,49].

### 2.6. Accession number

Raw metagenomic data and environmental genomes derived from binning processes were deposited in the Sequence Read Archive (SRA) under the bioproject accession number PRJNA602600 for raw data and PRJNA602601 for metagenome-assembled genomes.

## 3. Results

### 3.1. MAGs recovery

A total of ~3-4 million scaffolds were recovered from the soil metagenomes in each site. Even though all the sites underwent similar sequencing efforts (between 2.3 GB in 1640 and 1957 to 2.4 Gb in 1751), the youngest soil had the largest number of scaffolds with 499 sequences >50 kb compared to the oldest soil with only 12 scaffolds with a size >50 kb (Table 1). The youngest soil also had a larger N50 length of scaffolds and L50 compared to the other soils (Table 1). A total of 19 MAGs with a completeness of >70% and a contamination <5% (2 from 1640, 1 from 1751 and 16 from 1957) were retrieved and characterised.

**Table 1.** Summary report for the assembly quality assessment using MetaQuast.

| Statistics | 1640 | 1751 | 1957 |
| --- | --- | --- | --- |
| Number of scaffolds ($\geq 0$ bp) | 3631380 | 4047900 | 3138527 |
| Number of scaffolds ($\geq 500$ bp) | 1488437 | 1609020 | 1578887 |
| Number of scaffolds ($\geq 1000$ bp) | 326980 | 333885 | 447418 |
| Number of scaffolds ($\geq 25000$ bp) | 172 | 195 | 1551 |
| Number of scaffolds ($\geq 50000$ bp) | 12 | 40 | 499 |
| Total length ($\geq 0$ bp) | 2320437372 | 2460323814 | 2396069091 |
| Total length ($\geq 500$ bp) | 1385386471 | 1404094641 | 1717252425 |
| Total length ($\geq 1000$ bp) | 631179105 | 570404360 | 964085094 |
| Total length ($\geq 25000$ bp) | 5697241 | 8152100 | 82365310 |
| Total length ($\geq 50000$ bp) | 686375 | 2983958 | 46419897 |
| N50 | 914 | 850 | 700 |
| L50 | 391387 | 477193 | 348604 |

N50 - length such that scaffolds of this length or longer include half the bases of the assembly; L50 - number of scaffolds that are longer than, or equal to, the N50 length and therefore include half the bases of the assembly (https://www.ncbi.nlm.nih.gov/assembly/help/#globalstats)

### 3.2. MAG identification

MAGs were affiliated to the phyla Actinobacteria, Proteobacteria, Acidobacteria, Gemmatimonadetes, Chloroflexi, Firmicutes and Verrucomicrobia (Figure 2). In the oldest soil, two environmental genomes were retrieved related to Actinomycetales (Actinobacteria) and Rhodospirillales (Proteobacteria). The only MAG retrieved from the middle soil was related to Acidobacteria. MAGs binned from the youngest soil included seven assigned to Acidobacteria, one to Proteobacteria, three to Actinobacteria, one to the phylum Gemmatimonadetes, one to Verrucomicrobia and three to the phylum Chloroflexi (Figure 2).

**Figure 2.**
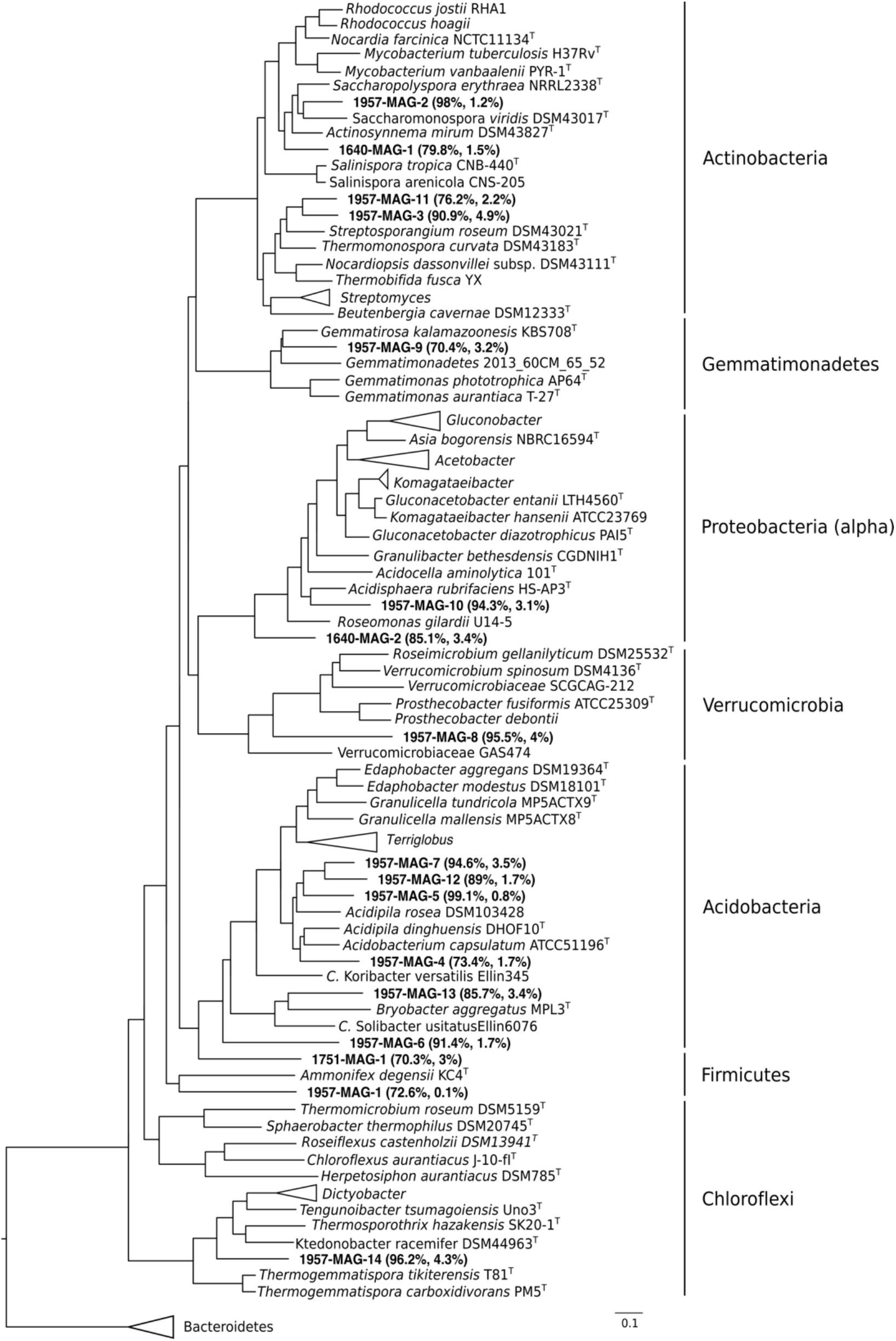
Phylogenomic tree of the bacterial genomic bins. The tree was built with PhyloSift against reference genomes downloaded from NCBI. MAGs are indicated in bold together with their respective completeness and contamination. FastTree confidence values of MAG branches are shown. The horizontal bar represents 10% sequence divergence.

The abundance of the MAGs in each site was calculated by using BLASTN (Figure 3). MAGs were more abundant from the soil they were recovered. MAGs with a total abundance >1% were found only in the young soil (1957). MAG 1957-2.1 (Ktedonobacteria, 1.21% ± 0.82), MAG 1957-5.1 (Actinomycetales, 1.02% ± 0.84), MAG 1957-13.1 (Verrucomicrobiales, 1.58% ± 1.1) and MAG 1957-16.1 (Acidobacteria, 1.14% ± 0.87) were the most abundant MAGs (Figure 3).

**Figure 3.**
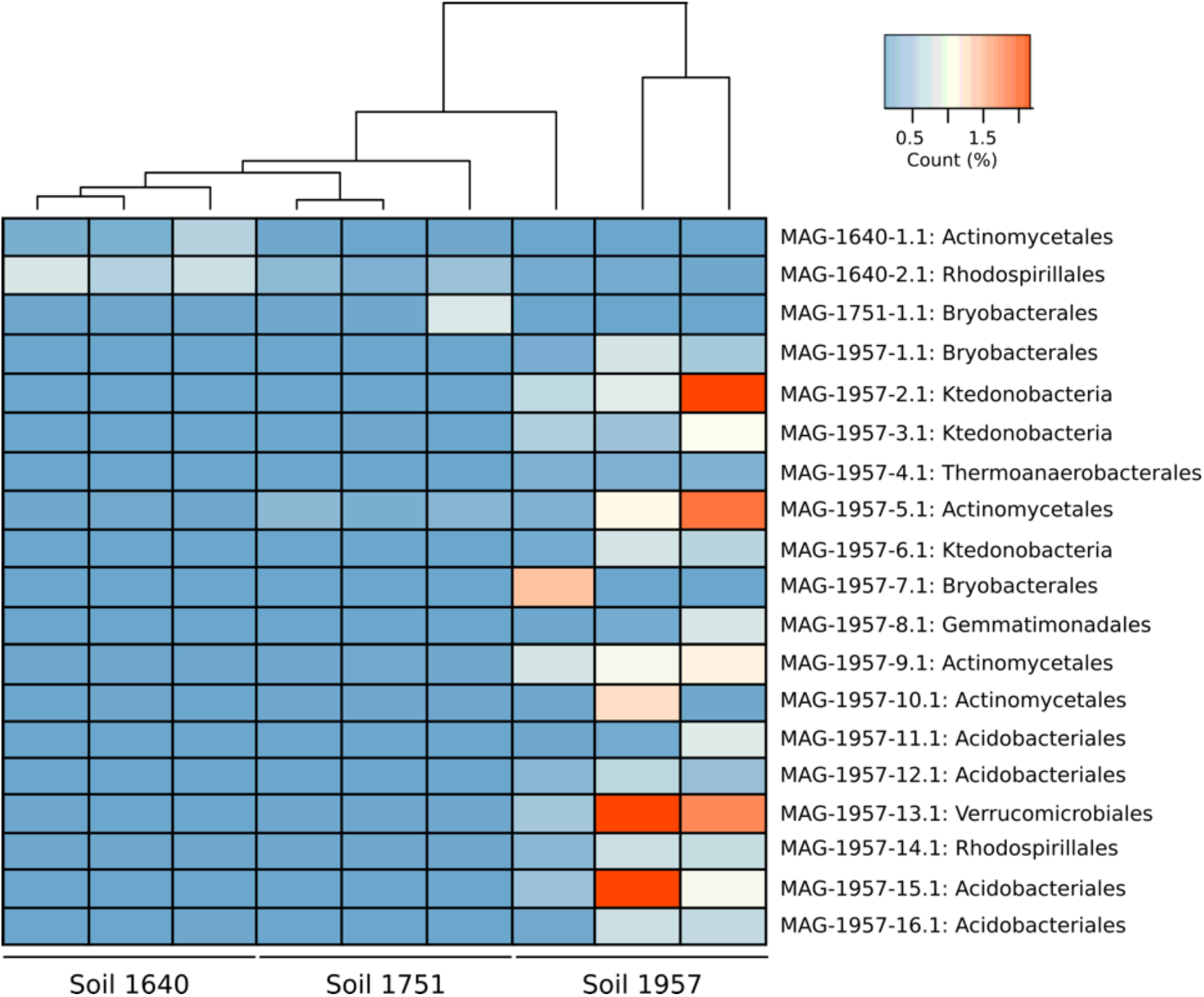
Heatmap representing the abundance of MAGs in each metagenome. The analysis was done by blast and only hits greater than 95% identity and e-value 1e-5 were used.

### 3.3. Metabolic characterisation of MAGs

Genes encoding enzymes involved in carbohydrate and energy metabolism, such as carbon fixation, sulfur metabolism, ATP synthesis and nitrogen metabolism, as well as terpenoid backbone biosynthesis were found in all the MAGs (Table 2). Other functions, including xenobiotic biodegradation, fatty acid metabolism, nucleotide metabolism, and vitamin metabolism, among others, were also found (Table S2).

**Table 2.**
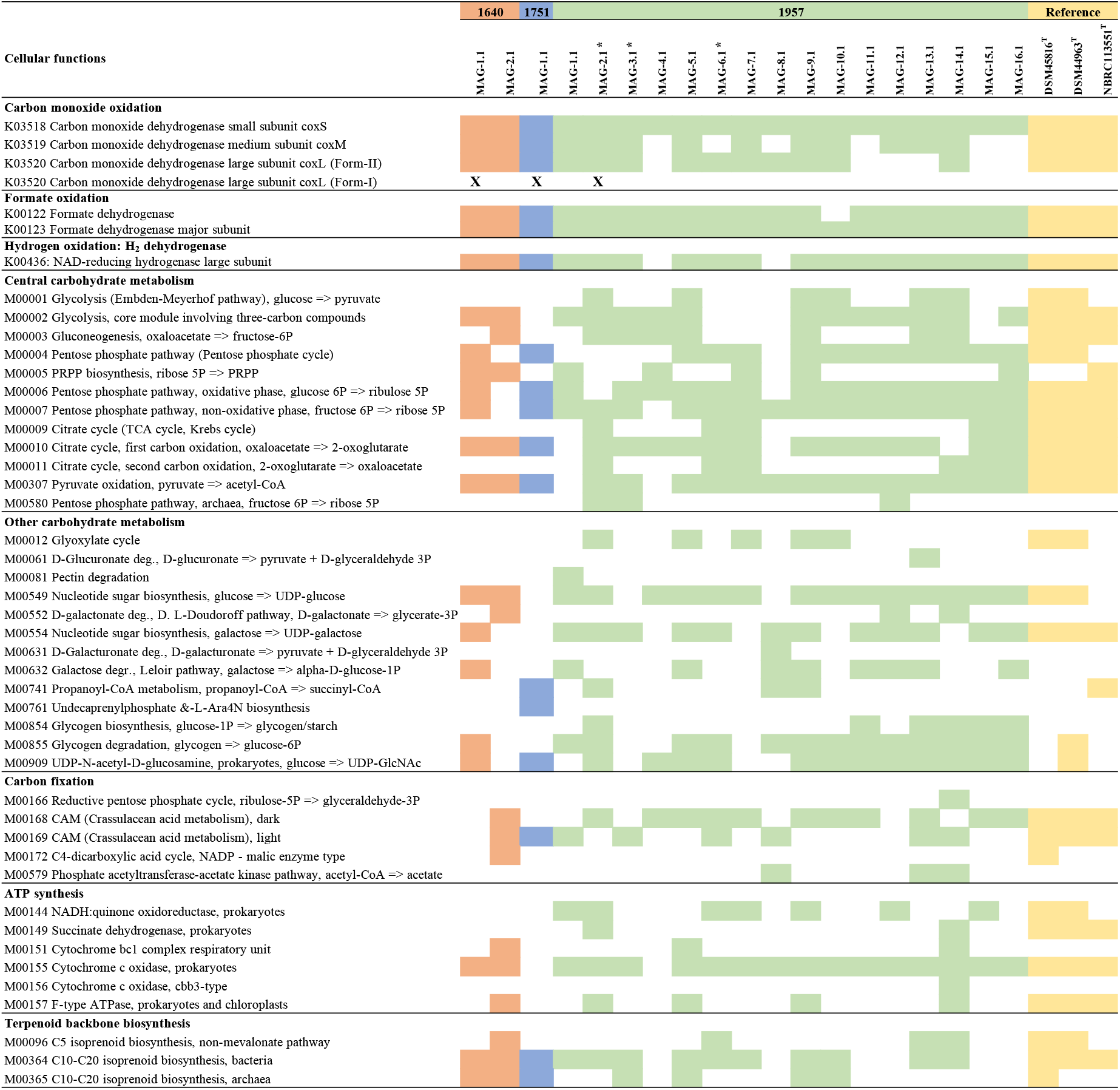
Summary table of complete cellular functions and other high-level features in the MAGs recovered form sites 1640, 1751 and 1957 retrieved from KEGG analysis. Reference genomes for Ketedonobacteria: DSM45816T [50], DSM44963T [51] and NBRC 113551T [52], (K: KEGG orthology; M: KEGG Mode). Asterisks indicate the MAGs isolated from the class Ktedonobacteria. “X” indicates MAGs containing genes encoding form I of the CoxL.

#### 3.3.1 Characterization of CODH and hydrogenase genes in MAGs

Three MAGs (MAG-1640-1.1, MAG-1751-1.1 and MAG-1957-2.1) encoded form I of the CO-dehydrogenase large subunit (*coxL*). These MAGs were each associated with a particular soil, with low abundance in the metagenomes of the other sites (Figure 4A). In addition to these three form-I *coxL* encoding MAGs, 15 other scaffolds from MAGs containing form II *coxL*-like genes were recovered (data not shown), but the function of form II CoxL is not yet known. The arrangement of genes encoding form I CODH in each of the MAGs is shown in Figure 4B. It should be noted that all of these three MAGs show the canonical arrangement for the three structural genes of CODH, that is the MSL genes. The genes encoding the [NiFe]-hydrogenase and its accessory proteins were only identified in MAG-1957-2.1, and instead only some of the accessory *hyp* genes were found in the other two MAGs (Figure 4B). A phylogenetic analysis of the form I *coxL* genes was performed, showing they are affiliated with Actinobacteria (MAG-1640-1.1), Nitrospirae *Candidatus* Manganitrophus noduliformans (MAG-1751-1.1) and Chloroflexi (MAG-1957-2.1) (Figure 5). This grouping is consistent with the results of PhyloSift (Figure 2), except for MAG-1751-1.1 where it was loosely associated with Acidobacteria (although with an amino-acid identity of only 40%) rather than Nitrospirae.

**Figure 4.**
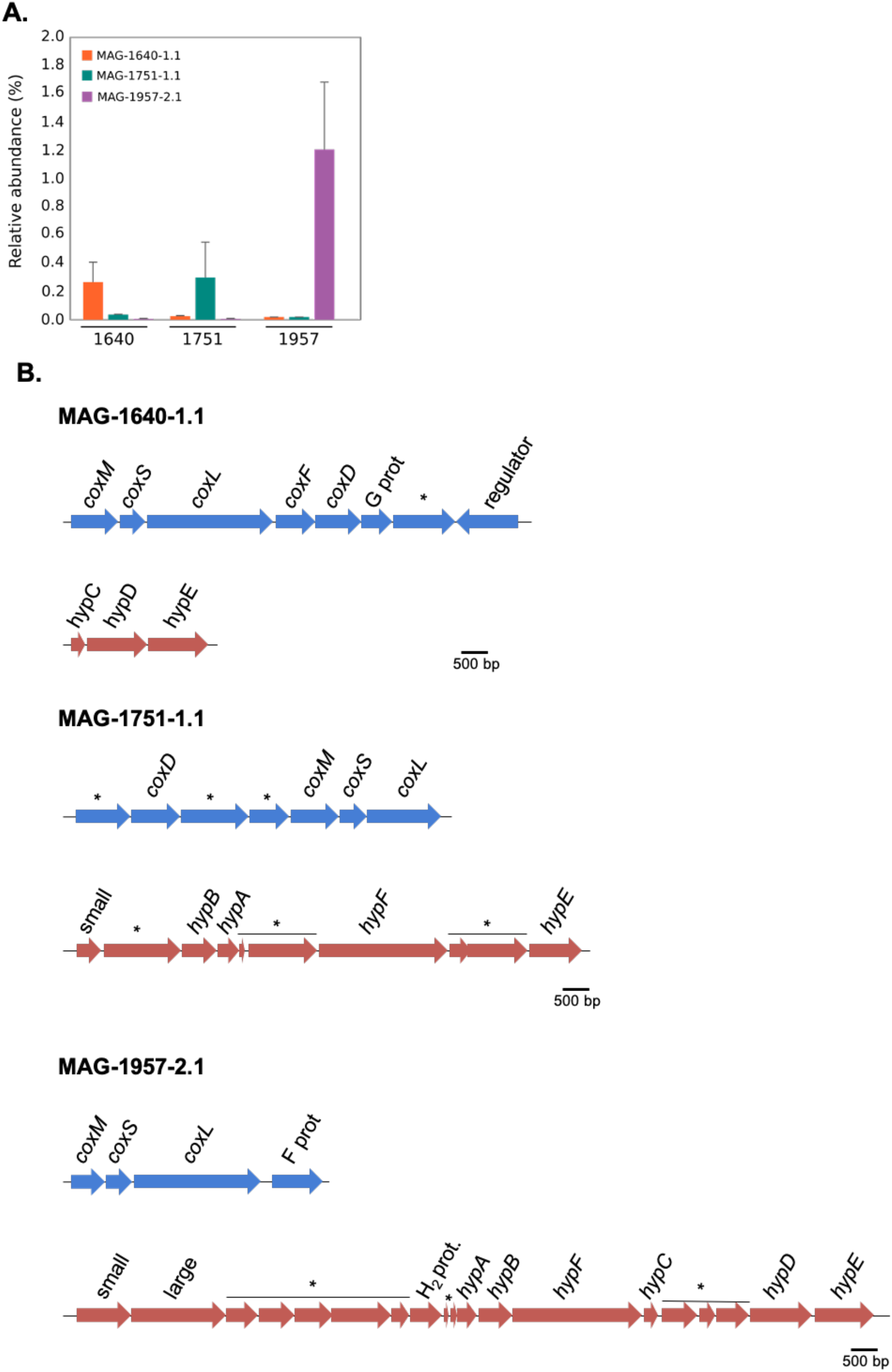
A. Relative abundance of MAG-1640-1.1, MAG-1751-1.1 and MAG-1957-2.1 in sites 1640, 1751, 1957. Bars indicate standard error of triplicates. B. Gene arrangement of the carbon monoxide dehydrogenase (CODH) and membrane-bound [NiFe]-hydrogenase in MAG-1640-1.1, MAG-1751-1.1 and MAG-1957-2.1 (* indicates hypothetical proteins; small indicates NiFe-hydrogenase small subunit and large indicates NiFe-hydrogenase large subunit).

**Figure 5.**
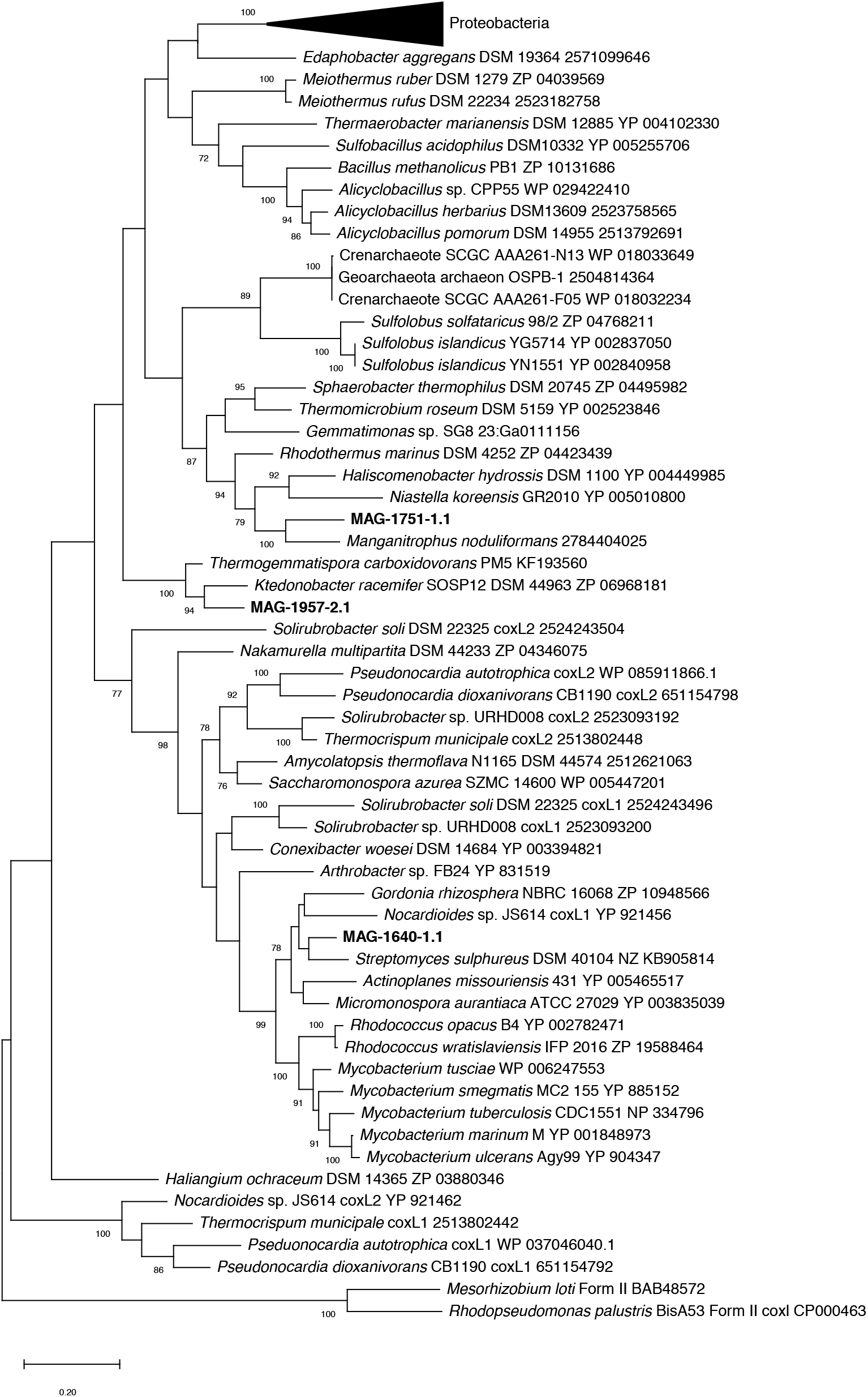
Phylogenetic tree of form I carbon monoxide dehydrogenase large subunit (CoxL) of metagenome-assembled genomes retrieved from Llaima volcano (MAG-1640-2.1, MAG-1751-1.1 and MAG-1957-2.1) against a reference sequences (with accession numbers included in the tree). The tree was drawn using the Maximum Likelihood method using MEGA X [48]. Bootstrap values (500 replications) are shown at the nodes. MAG-1751_1.1 is a partial sequence of the *coxL* gene.

#### 3.3.2 Complete metabolic characterization of Ktedonobacterales MAGs

Here we focused on the Ktedonobacterales MAGs because of their apparent importance in early soil formation. Three Ktedonobacterales MAGs were identified in the 1957 soil metagenomes, but were not found in the older soils (Figure 2 and 3). Two of the MAGs (MAGs 1957-2.1 and 1957-3.1), related/affiliated to the class Ktedonobacteria (phylum Chloroflexi), contained genes for the complete electron transport chain, citric acid metabolism, nitrogen metabolism, sulfur metabolism, several transporters, the complete gene set for carbon monoxide oxidation (CO dehydrogenase), herbicide degradation and degradation aromatics as well as the major subunit of the formate dehydrogenase, and also a hydrogenase. MAG 1957-6.1 (Ktedonobacteria) had very similar pathways as the other Chloroflexi MAGs, except a step for CO-oxidation and the electron transport chain were absent (Table 2, Table S2, Figure 6).

**Figure 6.**
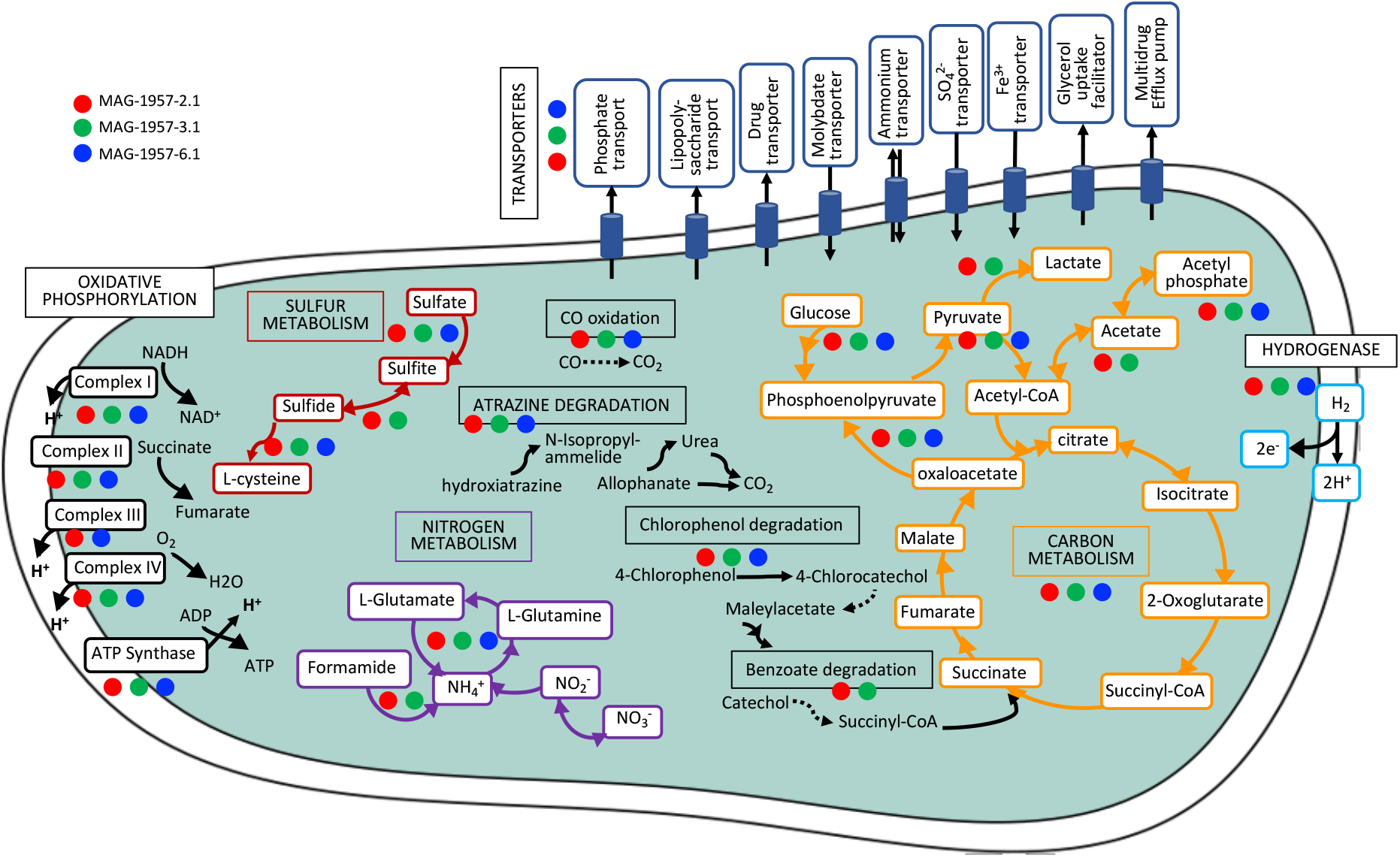
Metabolic reconstruction for some of the most important functions in the Ktedonobacteria MAGs isolated from the youngest soil using KEGG (More information in Table 2 and Table S2), dashed lines were used for incomplete pathways. Multi-arrows lines indicate several steps of a pathway.

## 4. Discussion

### 4.1. Characterisation of MAGs

In this study, a characterisation of metagenome-assembled genomes retrieved from Llaima volcano was performed. This study builds from a previous study [19] in which 16S rRNA gene amplicon-based sequences from those soils were analysed. The main objective of this study was to characterise genomes from those sites and to analyse the functions of the abundant but the poorly characterised Ktedonobacteria (phylum Chloroflexi) present at Llaima volcano. The relative abundance of the main phyla based on classification of scaffolds larger than 500 bp showed that microbial communities change as the soils age (Figure S1). This corroborates findings from a previous study [19]. For example, the relative abundance of Chloroflexi is higher in the younger soils (28% in the youngest soil to 7% in the oldest soil) and the opposite trend is observed for members of the phylum Proteobacteria, as their abundance increases as the soil ages (from 42% in the youngest soil to 59% in the oldest soil) (Figure S1). Except for those related to Firmicutes and Verrucomicrobia, and to a lesser extent Acidobacteria and Proteobacteria, the extracted environmental genomes had an amino acid identity > 50% with their closest reference genome (Table S3), which suggest that they belonged to those genera [53].

A total of 16 MAGs were recovered from the youngest soil (1957) (Figure 3). This soil is only partially vegetated (about 5%) by mosses and lichens. The microbial community in this area likely harbours populations able to grow as facultative chemolithoautotrophs or mixotrophs on carbon monoxide, hydrogen or methane. This high relative abundance of MAGs with genes for CO and hydrogen utilization in the youngest soils is consistent with reports by King and colleagues for Hawaiian and Japanese volcanic deposits (21-to 800-year old sites). For some of those sites, microbial community structure changed as the soil matured with members of the phylum Proteobacteria dominating vegetated sites while younger sites were enriched with Ktedonobacteria within the Chloroflexi and characterized by relatively high rates of atmospheric CO uptake [7,14,54].

MAGs were most abundant in the soil site from where they were retrieved (Figure 3). Relatively few MAGs were retrieved from the two older soils, which can be explained by the higher diversity in these soils and the decreased likelihood of recovering MAGs from groups such as Actinobacteria, Acidobacteria and Chloroflexi that were less common in them. In fact, several of the MAGs retrieved had a low relative abundance within the soils (Figure 3), which is consistent with their relative abundance of 16S rRNA genes in these soils [19]. Binning at the strain level remains a technical challenge [55], with the chances of retrieving MAGs at a given sequencing effort being reduced with increasing microdiversity (intra-population genetic diversity) and overall community diversity [56]. We previously reported that as the soil recovered and vegetation established, the microbial population appeared to enlarge and become more diverse [19], which explains the lower number of MAGS retrieved from more mature soil (1640 sample), compared to the younger sites (1957).

### 4.2. Metabolic characterisation of MAGs

The three MAGs containing form I *coxL* genes were found in an operon structure (Fig. 4B) typical of known CO oxidizers [41]. Form I *coxL* has been definitively associated with CO oxidation at high concentrations and also at sub-atmospheric levels [41]. Thus, even at low abundance, the presence of these *cox*-containing MAGs strongly suggests a capacity for atmospheric CO uptake at all the sites.

Most of the complete functions found from the Ktedonobacteria MAGs were also found in three reference genomes: *Ktedonobacter racemifer* DSM 44963 [51], *Thermogemmatispora carboxidivorans* PM5, isolated from a geothermal biofilm on Kilauea Volcano, Hawaii (USA) [50] and *Dictyobacter volcani* W12 [52]. According to our genomic analyses, all of these reference strains possess formate-, H_2_-, and CO-dehydrogenases as do the MAGs recovered in the present study. *Burkholderia* strains (phylum Proteobacteria) [57], members of the phylum Chloroflexi [14] and other members of the phyla Proteobacteria and Actinobacteria [58] have also been reported as CO-oxidisers in Hawaiian volcanic deposits. *coxL* genes encoding the large subunit of the CO dehydrogenase have been found in Proteobacteria species from Kilauea and Miyake-jima volcanoes [10,14,54].

The taxonomies of MAGs 1640-1.1 and 1957-2.1 were consistent for *coxL* (Figure 5) and phylogenomic analyses (Figure 2). In contrast, MAG-1751-1.1 clustered weakly with Acidobacteria based on genomic analysis (40% amino acid identity with a reference genome, see Table S3) but did not cluster with *Candidatus* Manganitrophus noduliformans as did the *coxL* sequence from this MAG. Several strains from the class Ktedonobacteria have been isolated from different environments (Table S4), but only *Dictyobacter vulcani* W12 [52] and *Thermogemmatispora carboxidivorans* PM5 [50] have been isolated from volcanic environments. So far, the class Ktedonobacteria contains only six genera and fifteen formally proposed species. Out of the 15 type strains, 11 genomes are available on RefSeq (Table S2). The order Ktedonobacterales contains the type strains *Ktedonobacter racemifer* SOSP1-21^T^ [51,59], *Dictyobacter aurantiacus* S-27^T^ [60], *Dictyobacter vulcani* W12^T^ [52], *Thermosporothrix hazakensis* SK20-1^T^ [61], *Thermosporothrix narukonensis* F4^T^ [62], *Ktedonosporobacter rubrisoli* SCAWS-G2^T^ [63], *Tengunoibacter tsumagoiensis* Uno3^T^, *Dictyobacter kobayashii* Uno11^T^ and *Dictyobacter alpinus* Uno16^T^ [64]. The order *Thermogemmatispora* contains the species *Thermogemmatispora aurantia* A1-2^T^ [65], *Thermogemmatispora argillosa* A3-2^T^ [65], *Thermogemmatispora onikobensis* NBRC 111776 (unpublished, RefSeq Nr NZ_BDGT00000000.1), *Thermogemmatispora onikobensis* ONI-1^T^ [66], *Thermogemmatispora foliorum* ONI-5^T^ [66] and *Thermogemmatispora carboxidivorans* PM5^T^ [50]. All of those genomes contain the complete gene set for carbon monoxide oxidation (CO dehydrogenase), as well as formate dehydrogenases, and a H_2_ dehydrogenases (Table S4). Our study particularly brings more insights into the role that early colonisers of this group from volcanic soils may have in the development of soils.

The large subunit of the NAD-reducing hydrogenase was also found in several MAGs (Table 2). Hydrogen metabolism has been shown to provide an additional energy source for some microorganisms and has been observed in bacteria and archaea [67]. Hydrogen dehydrogenases have also been found in members of the genus *Cupriavidus* (phylum Proteobacteria) from volcanic mudflow deposits in the Philippines suggesting their potential contribution to hydrogen uptake [68].

## 4. Conclusions

This study is further evidence that poorly characterised groups such as Ktedonobacteria, establish in remote volcanic sites and may use reduced gases for growth. Further studies are needed to demonstrate the activity of these pathways and their significance in volcanic deposits.

## Author Contributions

Conceptualization, M.H., R.C. and C.C.T.; sampling campaign, M.C.-F.; methodology, M.H. and B.V-G.; software, M.H.; validation, M.H., B.V-G., G.M.K., R.C. and C.C.T.; formal analysis, M.H.; investigation, M.H.; resources, M.H., R.C. and C.C.T.; data curation, M.H; writing—original draft preparation, M.H.; writing—review and editing, M.H., B.V-G., G.M.K., R.C. and C.C.T.; visualization, M.H.; supervision, R.C. and C.C.T.; project administration, M.H.; funding acquisition, M.H. All authors have read and agreed to the published version of the manuscript.

## Acknowledgments

We thank Mr. Stefan Mecke from the Thünen Institut für Biodiversität, Germany, for technical support. Part of the research presented in this paper was carried out on the High-Performance Computing Cluster supported by the Research and Specialist Computing Support Service (RSCSS) at the University of East Anglia; a special thanks to Mr. Mike Adams in RSCSS for technical support.

