## Supplementary Information for "Reconstructing genomes of carbon monoxide oxidisers in volcanic deposits including members of the class Ktedonobacteria"

**Supplementary figure legend:**
Figure S1. Relative abundance of main phyla based on classification of scaffolds >500 bp. Other phyla include members of Planctomycetes, Gemmatimonadetes, Cyanobacteria, Firmicutes, Armatimonadetes and Patescibacteria.

**Supplementary table legends:**
Table S1. List of strains used for the tree construction.
Table S2. Summary table of other complete cellular functions in the MAGs isolates form sites 1640, 1751 and 1957 retrieved from KEGG analysis. Reference genomes for Ketedonobacteria: DSM45816T [1], DSM44963$^T$ [2] and NBRC 113551$^T$ [3], (K: KEGG orthology; M: KEGG Mode). Asterisks indicate the MAGs isolated from the class Ktedonobacteria.
Table S3. Summary of the metagenome-assembled genomes (MAGs) isolated in the present study. Average amino-acid identity (AAI) was calculated by comparing the MAGs with their closest reference genomes identified based on the RAST results. The RefSeq accession number of the reference genomes are provided in parenthesis.
Table S4. Summary of enzymatic functions for CO-, H2-, and formate-oxidation in the Ktedonobacteria reference genomes. Carbon monoxide oxidation: K03518 carbon monoxide dehydrogenase small subunit coxS, K03519 carbon monoxide dehydrogenase medium subunit coxM, K03520 carbon monoxide dehydrogenase large subunit coxL. Formate oxidation: K00122 formate dehydrogenase, K00123 formate dehydrogenase major subunit. Hydrogen oxidation: $H_2$ dehydrogenase: K00436: NAD-reducing hydrogenase large subunit.

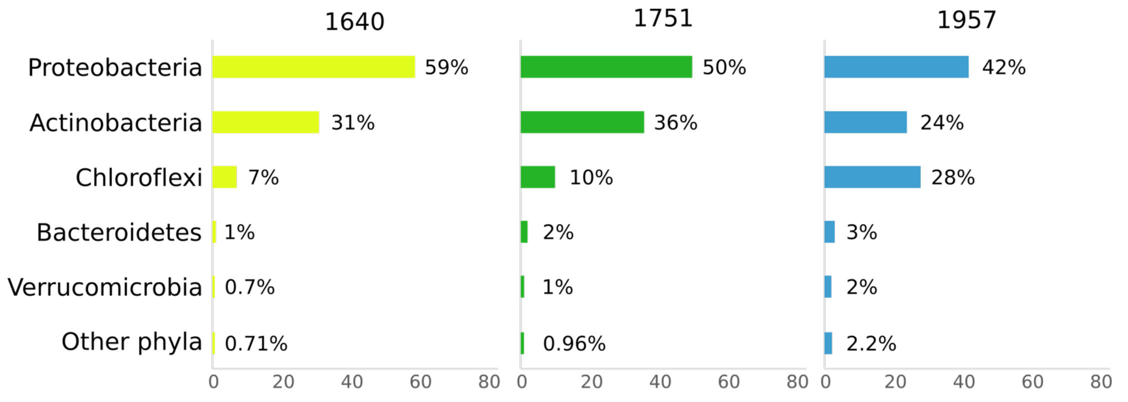


Figure S1. Relative abundance of main phyla based on classification of scaffolds >500 bp. Other phyla include members of Planctomycetes, Gemmatimonadetes, Cyanobacteria, Firmicutes, Armatimonadetes and Patescibacteria.

Table S1. List of strains used for the tree construction.

| Strain name | Type strain | NCBI Accession number |
|---|---|---|
| *Acetobacter aceti* TMW2.1153 | | NZ_CP014692.1 |
| *Acetobacter orleanensis* JCM 7639 | yes | NZ_BAMY01000269.1 |
| *Acetobacter pasteurianus* IFO 3283-01 | | NC_013209.1 |
| *Acetobacter syzygii* 9H-2 | | NZ_BAMZ01000094.1 |
| *Acidipila dinghuensis* DHOF10 | yes | NZ_SDMK01000001.1 |
| *Acidipila rosea* DSM 103428 | | NZ_SMGK01000001.1 |
| *Acidisphaera rubrifaciens* HS-AP3 | yes | NZ_BANB01001239.1 |
| *Acidobacterium capsulatum* ATCC 51196 | yes | NC_012483.1 |
| *Acidocella aminolytica* 101 | yes | NZ_FQVJ01000109.1 |
| *Actinosynnema mirum* DSM 43827 | yes | NC_013093.1 |
| *Ammonifex degensii* KC4 | yes | NC_013385.1 |
| *Asaia bogorensis* NBRC 16594 | yes | NZ_AP014690.1 |
| *Bacteroides fragilis* YCH46 | | NC_006347.1 |
| *Bacteroides ovatus* ATCC 8483 | yes | NZ_CP012938.1 |
| *Bacteroides uniformis* ATCC 8492 | yes | NZ_DS362249.1 |
| *Bacteroides vulgatus* ATCC 8482 | yes | NC_009614.1 |
| *Bacteroides dorei* CL03T12C01 | | NZ_CP011531.1 |
| *Beutenbergia cavernae* DSM 12333 | yes | NC_012669.1 |
| *Bryobacter aggregatus* MPL3 | yes | NZ_JNIF01000001.1 |
| *Candidatus* Koribacter versatilis Ellin345 | | NC_008009.1 |
| *Candidatus* Solibacter usitatus Ellin6076 | | NC_008536.1 |
| *Chloroflexus aurantiacus* J-10-fl | yes | NC_010175.1 |
| *Dictyobacter aurantiacus* S-27 | yes | NZ_BIFQ01000001.1 |
| *Dictyobacter alpinus* Uno16 | yes | NZ_BIFT01000001.1 |
| *Dictyobacter kobayashii* Uno11 | yes | NZ_BIFS01000001.1 |
| *Dictyobacter vulcani* W12 | yes | NZ_BKZW01000001.1 |
| *Edaphobacter aggregans* DSM 19364 | yes | NZ_JQKI01000001.1 |
| *Edaphobacter modestus* DSM 18101 | yes | NZ_SHKW01000001.1 |
| *Gemmatimonadetes* bacterium 2013_60CM_65_52 | | NGFI01000005.1 |
| *Gemmatimonas aurantiaca* T-27 | yes | NC_012489.1 |
| *Gemmatimonas phototrophica* AP64 | yes | NZ_CP011454.1 |
| *Gemmatirosa kalamazoonesis* KBS708 | yes | NZ_CP007128.1 |
| *Gluconacetobacter diazotrophicus* PAl 5 | yes | NC_011365.1 |
| *Gluconacetobacter entanii* LTH 4560 | yes | NZ_NKUF01000001.1 |
| *Gluconobacter kanchanaburiensis* NBRC 103587 | yes | NZ_BJVA01000001.1 |
| *Gluconobacter oxydans* H24 | | NC_019396.1 |
| *Gluconobacter roseus* LMG 1418 | | NZ_LHZP01000023.1 |
| *Gluconobacter sphaericus* NBRC 12467 | yes | NZ_BJMK01000001.1 |
| *Gluconobacter thailandicus* F149-1 | yes | NZ_BANH01000191.1 |
| *Gluconobacter wancherniae* NBRC 103581 | yes | NZ_BJUZ01000001.1 |
| *Granulicella mallensis* MP5ACTX8 | yes | NC_016631.1 |
| *Granulicella tundricola* MP5ACTX9 | yes | NC_015064.1 |
| *Herpetosiphon aurantiacus* DSM 785 | yes | CP000875.1 |

*continuation...*

| Strain name | Type strain | NCBI Accession number |
|---|---|---|
| *Komagataeibacter europaeus* SRCM101446 | | NZ_CP021467.1 |
| *Komagataeibacter hansenii* ATCC 23769 | | NZ_CM000920.1 |
| *Komagataeibacter oboediens* 174Bp2 | | NZ_CADT01000200.1 |
| *Komagataeibacter xylinus* E25 | | NZ_CP004360.1 |
| *Ktedonobacter racemifer* DSM 44963 | yes | NZ_ADVG01000010.1 |
| *Ktedonosporobacter rubrisoli* SCAWS-G2 | yes | NZ_CP035758.1 |
| *Mycobacterium tuberculosis* H37Rv | yes | NC_000962.3 |
| *Mycobacterium vanbaalenii* PYR-1 | yes | NC_008726.1 |
| *Nocardia farcinica* NCTC11134 | yes | NZ_LN868938.1 |
| *Nocardiopsis dassonvillei* DSM 43111 | yes | NC_014210.1 |
| *Prosthecobacter fusiformis* ATCC 25309 | yes | NZ_SOCA01000001.1 |
| *Prosthecobacter debontii* ATCC 700200 | yes | NZ_FUYE01000055.1 |
| *Rhodococcus jostii* RHA1 | | NC_008268.1 |
| *Roseiflexus castenholzii* DSM 13941 | yes | NC_009767.1 |
| *Roseimicrobium gellanilyticum* DSM 25532 | yes | NZ_QNRR01000001.1 |
| *Roseomonas gilardii* U14-5 | | NZ_CP015583.1 |
| *Saccharomonospora viridis* DSM 43017 | yes | NC_013159.1 |
| *Saccharopolyspora erythraea* NRRL2338 | yes | NC_009142.1 |
| *Salinispora arenicola* CNS-205 | | NC_009953.1 |
| *Salinispora tropica* CNB-440 | yes | NC_009380.1 |
| *Sphaerobacter thermophilus* DSM 20745 | yes | NC_013523.1 |
| *Streptomyces avermitilis* MA-4680 | yes | NC_003155.5 |
| *Streptomyces coelicolor* A3(2) | | NC_003888.3 |
| *Streptomyces griseus* subsp. griseus NBRC 13350 | | NC_010572.1 |
| *Streptomyces scabiei* 87.22 | | NC_013929.1 |
| *Streptosporangium roseum* DSM 43021 | yes | NC_013595.1 |
| *Tengunoibacter tsumagoiensis* Uno3 | yes | NZ_BIFR01000001.1 |
| *Terriglobus albidus* ORNL | | NZ_CP042806.1 |
| *Terriglobus roseus* DSM 18391 | yes | NC_018014.1 |
| *Terriglobus saanensis* SP1PR4 | yes | NC_014963.1 |
| *Terriglobus sp.* TAA 43 | | NZ_JUGR01000001.1 |
| *Thermobifida fusca* YX | | NC_007333.1 |
| *Thermogemmatispora carboxidivorans* PM5 | yes | NZ_JNIM01000001.1 |
| *Thermogemmatispora tikiterensis* T81 | yes | NZ_MCIF01000002.1 |
| *Thermogemmatispora aurantia* A1-2 | yes | NZ_BKZV01000001.1 |
| *Thermogemmatispora onikobensis* NBRC 111776 | yes | NZ_BDGT01000001.1 |
| *Thermomicrobium roseum* DSM 5159 | yes | NC_011959.1 |
| *Thermomonospora curvata* DSM 43183 | yes | NC_013510.1 |
| *Thermosporothrix hazakensis* SK20-1 | yes | NZ_BIFX01000001.1 |
| *Verrucomicrobium spinosum* DSM 4136 | yes | NZ_ABIZ01000001.1 |
| *Verrucomicrobium* sp. GAS474 | | LT629781 |

Table S2. Summary table of other complete cellular functions in the MAGs isolates form sites 1640, 1751 and 1957 retrieved from KEGG analysis. Reference genomes for Ketedonobacteria: DSM45816T [1], DSM44963$^{T}$ [2] and NBRC 113551$^{T}$ [3], (K: KEGG orthology; M: KEGG Mode). Asterisks indicate the MAGs isolated from the class Ktedonobacteria.

| | 1640 | | 1751 | 1957 | | | | | | | | | | | | | | | | Reference | | |
|---|---|---|---|---|---|---|---|---|---|---|---|---|---|---|---|---|---|---|---|---|---|---|
| **Function (M: KEGG Mode)** | MAG-1.1 | MAG-2.1 | MAG-1.1 | MAG-1.1 | MAG-2.1 * | MAG-3.1 * | MAG-4.1 | MAG-5.1 | MAG-6.1 * | MAG-7.1 | MAG-8.1 | MAG-9.1 | MAG-10.1 | MAG-11.1 | MAG-12.1 | MAG-13.1 | MAG-14.1 | MAG-15.1 | MAG-16.1 | DSM45816$^{T}$ | DSM44963$^{T}$ | NBRC113551$^{T}$ |
| **Fatty acid metabolism** | | | | | | | | | | | | | | | | | | | | | | |
| M00082 Fatty acid biosynthesis, initiation | | | | | | | | | | | | | | | | | | | | | | |
| M00083 Fatty acid biosynthesis, elongation | | | | | | | | | | | | | | | | | | | | | | |
| M00086 beta-Oxidation, acyl-CoA synthesis | | | | | | | | | | | | | | | | | | | | | | |
| M00087 beta-Oxidation | | | | | | | | | | | | | | | | | | | | | | |
| **Lipid metabolism** | | | | | | | | | | | | | | | | | | | | | | |
| M00091 Phosphatidylcholine (PC) biosynthesis, PE => PC | | | | | | | | | | | | | | | | | | | | | | |
| M00093 Phosphatidylethanolamine (PE) biosynthesis, PA => PS => PE | | | | | | | | | | | | | | | | | | | | | | |
| M00098 Acylglycerol degradation | | | | | | | | | | | | | | | | | | | | | | |
| **Purine metabolism** | | | | | | | | | | | | | | | | | | | | | | |
| M00048 Inosine monophosphate biosynthesis, PRPP + glutamine => IMP | | | | | | | | | | | | | | | | | | | | | | |
| M00049 Adenine ribonucleotide biosynthesis, IMP => ADP,ATP | | | | | | | | | | | | | | | | | | | | | | |
| M00050 Guanine ribonucleotide biosynthesis IMP => GDP,GTP | | | | | | | | | | | | | | | | | | | | | | |
| **Pyrimidine metabolism** | | | | | | | | | | | | | | | | | | | | | | |
| M00052 Pyrimidine ribonucleotide biosynthesis, UMP => UDP/UTP,CDP/CTP | | | | | | | | | | | | | | | | | | | | | | |
| M00046 Pyrimidine deg., uracil => beta-alanine, thymine => 3-aminoisobutanoate | | | | | | | | | | | | | | | | | | | | | | |
| **Serine and threonine metabolism** | | | | | | | | | | | | | | | | | | | | | | |
| M00018 Threonine biosynthesis, aspartate => homoserine => threonine | | | | | | | | | | | | | | | | | | | | | | |
| M00020 Serine biosynthesis, glycerate-3P => serine | | | | | | | | | | | | | | | | | | | | | | |
| M00555 Betaine biosynthesis, choline => betaine | | | | | | | | | | | | | | | | | | | | | | |
| **Cysteine and methionine metabolism** | | | | | | | | | | | | | | | | | | | | | | |
| M00017 Methionine biosynthesis, apartate => homoserine => methionine | | | | | | | | | | | | | | | | | | | | | | |
| M00021 Cysteine biosynthesis, serine => cysteine | | | | | | | | | | | | | | | | | | | | | | |
| M00035 Methionine degradation | | | | | | | | | | | | | | | | | | | | | | |
| M00338 Cysteine biosynthesis, homocysteine + serine => cysteine | | | | | | | | | | | | | | | | | | | | | | |
| **Branched-chain amino acid metabolism** | | | | | | | | | | | | | | | | | | | | | | |
| M00019 Val/isoleucine biosynthesis, pyruvate => val/2-oxobutanoate => isoleucine | | | | | | | | | | | | | | | | | | | | | | |
| M00432 Leucine biosynthesis, 2-oxoisovalerate => 2-oxoisocaproate | | | | | | | | | | | | | | | | | | | | | | |
| M00535 Isoleucine biosynthesis, pyruvate => 2-oxobutanoate | | | | | | | | | | | | | | | | | | | | | | |
| M00570 Isoleucine biosynthesis, threonine => 2-oxobutanoate => isoleucine | | | | | | | | | | | | | | | | | | | | | | |
| **Arginine and proline metabolism** | | | | | | | | | | | | | | | | | | | | | | |
| M00015 Proline biosynthesis, glutamate => proline | | | | | | | | | | | | | | | | | | | | | | |
| M00028 Ornithine biosynthesis, glutamate => ornithine | | | | | | | | | | | | | | | | | | | | | | |
| M00844 Arginine biosynthesis, ornithine => arginine | | | | | | | | | | | | | | | | | | | | | | |
| **Polyamine biosynthesis** | | | | | | | | | | | | | | | | | | | | | | |
| M00134 Polyamine biosynthesis, arginine => ornithine => putrescine | | | | | | | | | | | | | | | | | | | | | | |
| M00135 GABA biosynthesis, eukaryotes, putrescine => GABA | | | | | | | | | | | | | | | | | | | | | | |
| **Aromatic amino acid metabolism** | | | | | | | | | | | | | | | | | | | | | | |
| M00022 Shikimate pathway, phosphoenolpyruvate + erythrose-4P => chorismate | | | | | | | | | | | | | | | | | | | | | | |
| M00023 Tryptophan biosynthesis, chorismate => tryptophan | | | | | | | | | | | | | | | | | | | | | | |
| **Histidine metabolism** | | | | | | | | | | | | | | | | | | | | | | |
| M00026 Histidine biosynthesis, PRPP => histidine | | | | | | | | | | | | | | | | | | | | | | |
| M00045 Histidine degradation, histidine => N-formiminoglutamate => glutamate | | | | | | | | | | | | | | | | | | | | | | |
| **Lysine metabolism** | | | | | | | | | | | | | | | | | | | | | | |
| M00016 Lysine biosynthesis, succinyl-DAP pathway, aspartate => lysine | | | | | | | | | | | | | | | | | | | | | | |
| M00527 Lysine biosynthesis, DAP aminotransferase pathway, aspartate => lysine | | | | | | | | | | | | | | | | | | | | | | |
| **Other amino acid metabolism** | | | | | | | | | | | | | | | | | | | | | | |
| M00027 GABA (gamma-Aminobutyrate) shunt | | | | | | | | | | | | | | | | | | | | | | |
| M00118 Glutathione biosynthesis, glutamate => glutathione | | | | | | | | | | | | | | | | | | | | | | |
| **Lipopolysaccharide metabolism** | | | | | | | | | | | | | | | | | | | | | | |
| M00063 CMP-KDO biosynthesis | | | | | | | | | | | | | | | | | | | | | | |
| M00866 KDO2-lipid A biosynthesis, Raetz pathway, non-LpxL-LpxM type | | | | | | | | | | | | | | | | | | | | | | |
| **Cofactor and vitamin metabolism** | | | | | | | | | | | | | | | | | | | | | | |
| M00115 NAD biosynthesis, aspartate => NAD | | | | | | | | | | | | | | | | | | | | | | |
| M00119 Pantothenate biosynthesis, valine/L-aspartate => pantothenate | | | | | | | | | | | | | | | | | | | | | | |
| M00120 Coenzyme A biosynthesis, pantothenate => CoA | | | | | | | | | | | | | | | | | | | | | | |
| M00121 Heme biosynthesis, plants and bacteria, glutamate => heme | | | | | | | | | | | | | | | | | | | | | | |
| M00123 Biotin biosynthesis, pimeloyl-ACP/CoA => biotin | | | | | | | | | | | | | | | | | | | | | | |
| M00140 C1-unit interconversion, prokaryotes | | | | | | | | | | | | | | | | | | | | | | |
| M00577 Biotin biosynthesis, BioW pathway, pimelate => pimeloyl-CoA => biotin | | | | | | | | | | | | | | | | | | | | | | |
| M00846 Siroheme biosynthesis, glutamate => siroheme | | | | | | | | | | | | | | | | | | | | | | |
| M00868 Heme biosynthesis, animals and fungi, glycine => heme | | | | | | | | | | | | | | | | | | | | | | |
| M00880 Molybdenum cofactor biosynthesis, GTP => molybdenum cofactor | | | | | | | | | | | | | | | | | | | | | | |
| **Sulfur metabolism** | | | | | | | | | | | | | | | | | | | | | | |
| M00176 Assimilatory sulfate reduction, sulfate => H2S | | | | | | | | | | | | | | | | | | | | | | |
| **Nitrogen metabolism** | | | | | | | | | | | | | | | | | | | | | | |
| M00531 Assimilatory nitrate reduction, nitrate => ammonia | | | | | | | | | | | | | | | | | | | | | | |
| **Polyketide sugar unit biosynthesis** | | | | | | | | | | | | | | | | | | | | | | |
| M00793 dTDP-L-rhamnose biosynthesis | | | | | | | | | | | | | | | | | | | | | | |
| **Aromatics degradation** | | | | | | | | | | | | | | | | | | | | | | |
| M00551 Benzoate deg., benzoate => catechol/methylbenzoate => methylcatechol | | | | | | | | | | | | | | | | | | | | | | |
| **Metabolic capacity** | | | | | | | | | | | | | | | | | | | | | | |
| M00616 Sulfate-sulfur assimilation | | | | | | | | | | | | | | | | | | | | | | |
| M00615 Nitrate assimilation | | | | | | | | | | | | | | | | | | | | | | |

Table S3. Summary of the metagenome-assembled genomes (MAGs) isolated in the present study. Average amino-acid identity (AAI) was calculated by comparing the MAGs with their closest reference genomes identified based on the RAST results. The RefSeq accession number of the reference genomes are provided in parenthesis.

| MAG | Completeness (%) | Contamination (%) | GC (%) | N50 | Size (bp) | Phyla | Class/Order | Reference genome | AAI (%) |
|---|---|---|---|---|---|---|---|---|---|
| MAG-1640-1.1 | 80.57 | 1.536 | 69.1 | 8867 | 5073303 | Actinobacteria | Actinomycetales | *Actinosynnema mirum* DSM 43827$^{T}$ (NC_013093.1) | 58.65 |
| MAG-1640-2.1 | 85.36 | 3.738 | 66.8 | 11294 | 5751495 | Proteobacteria | Rhodospirillales | *Roseomonas gilardii* U14-5 (NZ_CP015583.1) | 49.1 |
| MAG-1751-1.1 | 70.3 | 3.1 | 70.8 | 7414 | 4752807 | Acidobacteria | Bryobacterales | *Candidatus* Koribacter versatilis Ellin345 (NC_008009.1) | 40.96 |
| MAG-1957-1.1 | 85.8 | 3.48 | 58.3 | 67512 | 4938955 | Acidobacteria | Bryobacterales | *Bryobacter aggregatus* MPL3$^{T}$ (NZ_JNIF01000001.1) | 49.77 |
| MAG-1957-2.1 | 96.2 | 3.96 | 53.4 | 119701 | 6327585 | Chloroflexi | Ktedonobacteria | *Ktedonosporobacter rubrisoli* SCAWS-G2$^{T}$ (NZ_CP035758.1 ) | 59.01 |
| MAG-1957-3.1 | 72.22 | 3.47 | 51.7 | 13375 | 6142618 | Chloroflexi | Ktedonobacteria | *Dictyobacter vulcani* W12$^{T}$ (NZ_BKZW01000001.1) | 63.43 |
| MAG-1957-4.1 | 72.64 | 0.1 | 67.2 | 5165 | 3007516 | Firmicutes | Thermoanaerobacterales | *Ammonifex degensii* KC4$^{T}$ (NC_013385.1) | 41.36 |
| MAG-1957-5.1 | 98.05 | 0.9 | 71.4 | 79391 | 5837494 | Actinobacteria | Actinomycetales | *Actinosynnema mirum* DSM 43827$^{T}$ (NC_013093.1) | 55.92 |
| MAG-1957-6.1 | 80.36 | 1.98 | 54.8 | 22264 | 5685492 | Chloroflexi | Ktedonobacteria | *Ktedonosporobacter rubrisoli* SCAWS-G2$^{T}$ (NZ_CP035758.1 ) | 58.66 |
| MAG-1957-7.1 | 94.02 | 1.71 | 58.9 | 137749 | 4659534 | Acidobacteria | Bryobacterales | *Candidatus* Koribacter versatilis Ellin345 (NC_008009.1) | 47.98 |
| MAG-1957-8.1 | 70.49 | 3.3 | 72.5 | 10863 | 4227183 | Gemmatimonadetes | Gemmatimonadales | *Gemmatirosa kalamazoonesis* KBS708$^{T}$ (NZ_CP007128.1) | 60.46 |
| MAG-1957-9.1 | 90.97 | 4.94 | 71.3 | 26163 | 6509599 | Actinobacteria | Actinomycetales | *Streptosporangium roseum* DSM 43021$^{T}$ (NC_013595.1) | 52.26 |
| MAG-1957-10.1 | 72.61 | 1.07 | 71.4 | 17599 | 4220629 | Actinobacteria | Actinomycetales | *Streptosporangium roseum* DSM 43021$^{T}$ (NC_013595.1) | 52.93 |
| MAG-1957-11.1 | 89.03 | 1.72 | 60.9 | 88061 | 3399414 | Acidobacteria | Acidobacteriales | *Acidipila rosea* DSM 103428 (NZ_SMGK01000001.1) | 54.36 |
| MAG-1957-12.1 | 73.41 | 1.72 | 58.9 | 22191 | 4793286 | Acidobacteria | Acidobacteriales | *Acidobacterium capsulatum* ATCC 51196$^{T}$ (NC_012483.1) | 57.18 |
| MAG-1957-13.1 | 95.6 | 4.09 | 60.5 | 13562 | 5255071 | Verrucomicrobia | Verrucomicrobiales | *Prosthecobacter debontii* ATCC 700200$^{T}$ (NZ_FUYE01000055.1) | 44.57 |
| MAG-1957-14.1 | 96.79 | 3.19 | 61.9 | 29422 | 6536060 | Proteobacteria | Rhodospirillales | *Acidisphaera rubrifaciens* HS-AP3$^{T}$ (NZ_BANB01001239.1) | 61.34 |
| MAG-1957-15.1 | 94.61 | 3.54 | 61.5 | 71986 | 3914044 | Acidobacteria | Acidobacteriales | *Acidipila rosea* DSM 103428 (NZ_SMGK01000001.1) | 57.33 |
| MAG-1957-16.1 | 99.14 | 0.86 | 65.4 | 109666 | 3669710 | Acidobacteria | Acidobacteriales | *Acidipila rosea* DSM 103428 (NZ_SMGK01000001.1) | 57.44 |

Table S4. Summary of enzymatic functions for CO-, H2-, and formate-oxidation in the Ktedonobacteria reference genomes. Carbon monoxide oxidation: K03518 carbon monoxide dehydrogenase small subunit coxS, K03519 carbon monoxide dehydrogenase medium subunit coxM, K03520 carbon monoxide dehydrogenase large subunit coxL. Formate oxidation: K00122 formate dehydrogenase, K00123 formate dehydrogenase major subunit. Hydrogen oxidation: $H_2$ dehydrogenase: K00436: NAD-reducing hydrogenase large subunit.

| Species | Strain | Isolation source | RefSeq | Collection | Reference | K03518 | K03519 | K03520 | K00122 | K00123 | K00436 |
|---|---|---|---|---|---|---|---|---|---|---|---|
| *Ktedonobacter racemifer* | SOSP1-21$^T$ | European soils | NZ_ADVG00000000.1 | DSM 44963 | [2] | x | x | x | x | x | x |
| *Dictyobacter aurantiacus* | S-27$^T$ | paddy soils | NZ_BIFQ00000000.1 | ASM396751v1 | [4] | x | x | x | x | x | x |
| *Dictyobacter vulcani* | W12$^T$ | volcanic soils | NZ_BKZW00000000.1 | NBRC113551T | [3] | x | x | x | x | x | x |
| *Thermosporothrix hazakensis* | SK20-1$^T$ | ripe compost | NZ_BIFX00000000.1 | ATCC BAA-1881 | [5] | x | x | x | x | x | x |
| *Ktedonosporobacter rubrisoli* | SCAWS-G2$^T$ | red soils | GCF_004208415.1 | DSM:105258 | [6] | x | x | x | x | x | x |
| *Tengunoibacter tsumagoiensis* | Uno3$^T$ | soil-like granular mass | NZ_BIFR00000000.1 | ASM396753v1 | [7] | x | x | x | x | x | x |
| *Dictyobacter kobayashii* | Uno11$^T$ | soil-like granular mass | NZ_BIFS00000000.1; | ASM396755v1 | [7] | x | x | x | x | x | x |
| *Dictyobacter alpinus* | Uno16$^T$ | soil-like granular mass | NZ_BIFT00000000.1 | ASM396757v1 | [7] | x | x | x | x | x | x |
| *Thermogemmatispora aurantia* | A1-2$^T$ | geothermal soil | NZ_BKZV00000000.1 | ASM897428v1 | [8] | x | x | x | x | x | x |
| *Thermogemmatispora onikobensis* | NBRC 111776 | geothermal soil | NZ_BDGT00000000.1 | ASM174828v1 | unpublished | x | x | x | x | x | x |
| *Thermogemmatispora carboxidivorans* | PM5$^T$ | biofilm | NZ_JNIM00000000.1; | DSM45816T | [1] | x | x | x | x | x | x |
